# Functional rather than anatomic connectivity predicts seizure propagation in a multi-node model of focal neocortical epilepsy

**DOI:** 10.1101/2025.08.26.672408

**Authors:** James E. Niemeyer, Fengrui Zhan, Carmen Pons, Theodore H. Schwartz

## Abstract

Seizures propagate through the brain either locally or via widespread networks of anatomically and functionally connected nodes. These sites can be manipulated in the surgical treatment of human patients through ablation or stimulation. However, we still lack a full understanding of how seizures, ablation, and stimulation recruit or alter recruitment of these distant sites. Here, we apply widefield calcium imaging in a non-anesthetized rodent multi-nodal bilateral neocortical network model of focal epilepsy to examine excitatory and inhibitory cell recruitment. When we initiate seizures in somatosensory cortex (S1), they preferentially spread to an ipsilateral node in frontal cortex (M2) rather than across the corpus callosum to contralateral mirror somatosensory cortex. On the other hand, seizures rapidly spread across the corpus callosum in regions that connect M2 with its mirror M2 focus, indicating that this frontal region acts an amplifier for secondary generalization. Accordingly, ablation of M2 radically altered seizure propagation. Electrical stimulation of S1 revealed that S1 preferentially recruits excitatory cells in ipsilateral M2 but inhibitory cells in contralateral S1, which may explain the preferred propagation pathway. We also observed that the stimulation frequency can differentially determine the response of excitatory versus inhibitory neurons. Altogether, our findings suggest that seizures do not propagate homogeneously through anatomically connected nodes but are preferentially “pulled” to specific locations by excitatory/inhibitory balance. Thus, functional connectivity rather than anatomic connectivity will be more predictive of ictal spread, and more informative for ablative and stimulation-based therapeutics.

**Significance:** Understanding how seizures spread outward from one brain region to another is critical to informing new therapies for epilepsy patients. We used widefield calcium imaging and electrical stimulation to examine seizure propagation patterns in a bilateral brain network in non-anesthetized animals. We found that seizures unexpectedly preferred certain pathways over others, despite strong anatomical connections, and that these preferences were dictated by the type of cell (excitatory or inhibitory) recruited in distant nodes. We were able to reproduce these electrical imbalances with focal stimulation at varying frequencies. Our findings highlight that anatomy by itself is insufficient to fully identify seizure networks and suggests that specific neurostimulation parameters will confer different effects on distant brain regions.

**Key Points:**

- Functional connectivity rather than anatomic connectivity determines ictal propagation
- Contralateral seizure propagation from S1 favors cross callosal spread in via motor, rather than somatosensory pathways.
- High frequency electrical stimulation differentially affects excitatory and inhibitory recruitment across a neocortical network
- Low frequency electrical stimulation emulates seizure-like activity propagation across a network
- Connectome data may be insufficient to identify critical nodes and pathways of a seizure network

## Introduction

Epilepsy afflicts about 1% of the population^1^ and exhibits drug resistance in over 30% of patients^2^. For these patients, the remaining surgical treatment options include disconnection, ablation, and neurostimulation. Epilepsy is increasingly considered to be a disorder of brain networks, in which multiple regions can be involved in the onset and spread of seizures as well as the long-term maintenance of epilepsy^3-5^. Various studies have highlighted the importance of network considerations in epilepsy resection and targeted neurostimulation therapies^6-12^. In line with this thinking, scientists employ graph theory methods to identify features of human brain network nodes (anatomical regions), such as effective connectivity^13^, hubness^14^, and centrality^15^ that correlate with identification of the seizure onset zone or extra-focal nodes involved in epilepsy that may be amenable to control^16, 17^. The cumulative results of this work strongly suggest that network neuroscience can provide improvements to surgical interventions in epilepsy^6, 7, 18^. However, despite this promising vision, we still lack a complete understanding of seizure dynamics through brain networks, partly due to a lack of robust animal models of multi-nodal seizure networks.

Seizures initiate from the spread of synchronous and asynchronous coordinated neuronal activity. This propagation has historically been characterized as contiguous spread – like a countertop spill – as in the “Jacksonian march”^19, 20^. Ictal spread can also be non-contiguous, or saltatory, when seizures hop from one computational column or module to the next^21-23^, or to more distant nodes through long-range white matter connections^24-30^. Some studies have even identified a role for distant brain sites in the *suppression* of seizures during interictal periods^6, 31^. While there may exist variability between patients, and across *in vivo* and *in vitro* animal models, it is expected that a comprehensive understanding of how seizures spread, particularly across bilateral brain regions, will improve surgical targeting of key nodes within an epileptic network. For example, ablation of an amplifying node may suppress seizure spread while ablation of a suppressing node may have the opposite effect.

Another component of seizures that remains poorly understood is the role of excitatory and inhibitory cellular activity during seizure spread. Two nodes in a network can be anatomically connected, but this connection may be functionally excitatory or inhibitory, depending on what cell type is activated at the distant synapse and the nature of the neurotransmitters released. This cell-specific connectivity is also of major relevance in therapeutic neurostimulation. Standard neurostimulation parameters typically employ high frequencies^18, 32^ due to findings that these frequencies are associated with suppressive effects^33-35^ or desynchronization^36-38^. By identifying stimulation parameters that could preferentially recruit or inhibit certain cell types (e.g., inhibitory or excitatory cells, respectively), researchers may enhance the efficacy of neurostimulatory treatments in epilepsy. Recent studies with cellular resolution imaging during electrical stimulation show that stimulation can differentially affect excitatory and inhibitory cells^39, 40^. However, little is known about how a focal electrical stimulation exerts effects across a widespread brain network.

Here we used widefield calcium imaging and electrophysiology to identify a multi-node seizure network in non-anesthetized mice. By imaging excitatory and inhibitory cell types, we examined the recruitment of different cells during ongoing seizure activity across a widespread network of monosynaptically connected nodes. Our findings demonstrate that seizures recruit preferential pathways of propagation, favoring certain nodes over others, depending on the E/I balance within that node, and that focal stimulation and ablation can selectively recapitulate and reroute seizure propagation. Our study provides new insights into the inhomogeneities in seizure propagation and the impact of long-range synaptic connections, excitatory/inhibitory (E/I) balance, and frequency-dependent cell-specific recruitment.

## Results

We identified a bilateral cortical anatomical network based on known white matter connections between primary somatosensory cortex (S1), ipsilateral secondary motor cortex (M2), and contralateral S1 and M2 **(Fig. 1A)**^41, 42^. Injection of 4-Aminopyridine (4-AP) injection into S1 resulted in electrographic seizure propagation across the 4-node bilateral network **(Fig. 1B-C**). The electrographic profile of this seizure showed a propagation pattern of earliest activity at the injection site in ipsilateral (i)S1, followed by ipsilateral (i)M2, followed by contralateral (c)M2, and finally recruitment of contralateral (c)S1. This pattern suggested that seizure propagation in the S1-M2 network did not immediately recruit cS1 but rather showed faster recruitment of the iS1-iM2 pathway. Most surprisingly, however, was that cM2, a node that is not monosynaptically connected to iS1, was recruited several seconds earlier than cS1. This data suggested that the S1-initiated seizure spreads preferentially along certain white matter tracts over others; namely, we found preferential spread anteriorly across the frontal motor component of the corpus callosum (iM2 to cM2) rather than across the more posterior corpus callosum which carries sensory information (iS1 to cS1).

**Figure 1.**
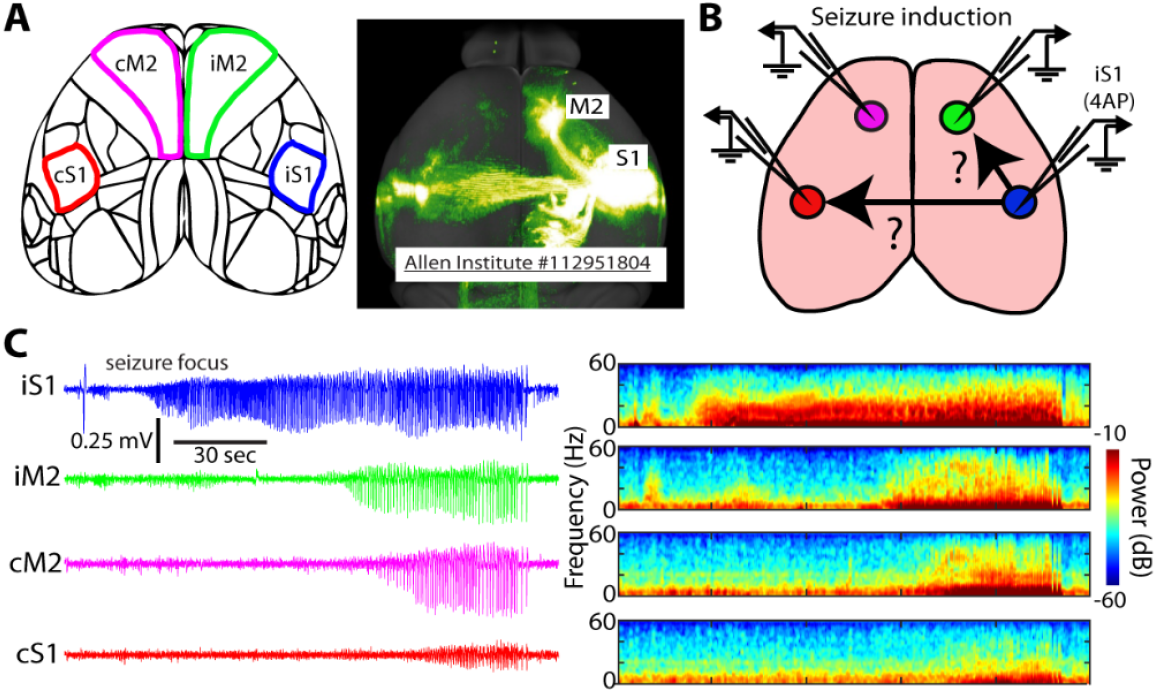
Network investigation. A) Allen institute mapping of S1 projections in the dorsal neocortex shows strong output to ipsilateral M2 and contralateral S1 (https://connectivity.brain-map.org/projection/experiment/112951804). B) Schematic depiction of a pilot experiment examining potential propagation pathways between monosynaptically connected nodes. Separate LFP electrodes are placed in each of the 4 network nodes (iS1, iM2, cM2, cS1). 4-AP is injected into iS1. C) Seizure activity spreads sequentially from iS1 to iM2 to cM2 to cS1; Left: LFP, Right: Power spectrogram).

To increase our spatial sampling, we next employed widefield calcium imaging of bilateral dorsal neocortex encompassing all four nodes during focal seizures initiating by 4-AP in iS1 in GCaMP6f-expressing mice (**Fig. 2**). We applied a custom line-length mapping method^43^ to create a propagation map that describes seizure initiation and propagation patterns, which shows high agreement with the local field potential line-length (**Supplemental Fig. 1**).

**Figure 2.**
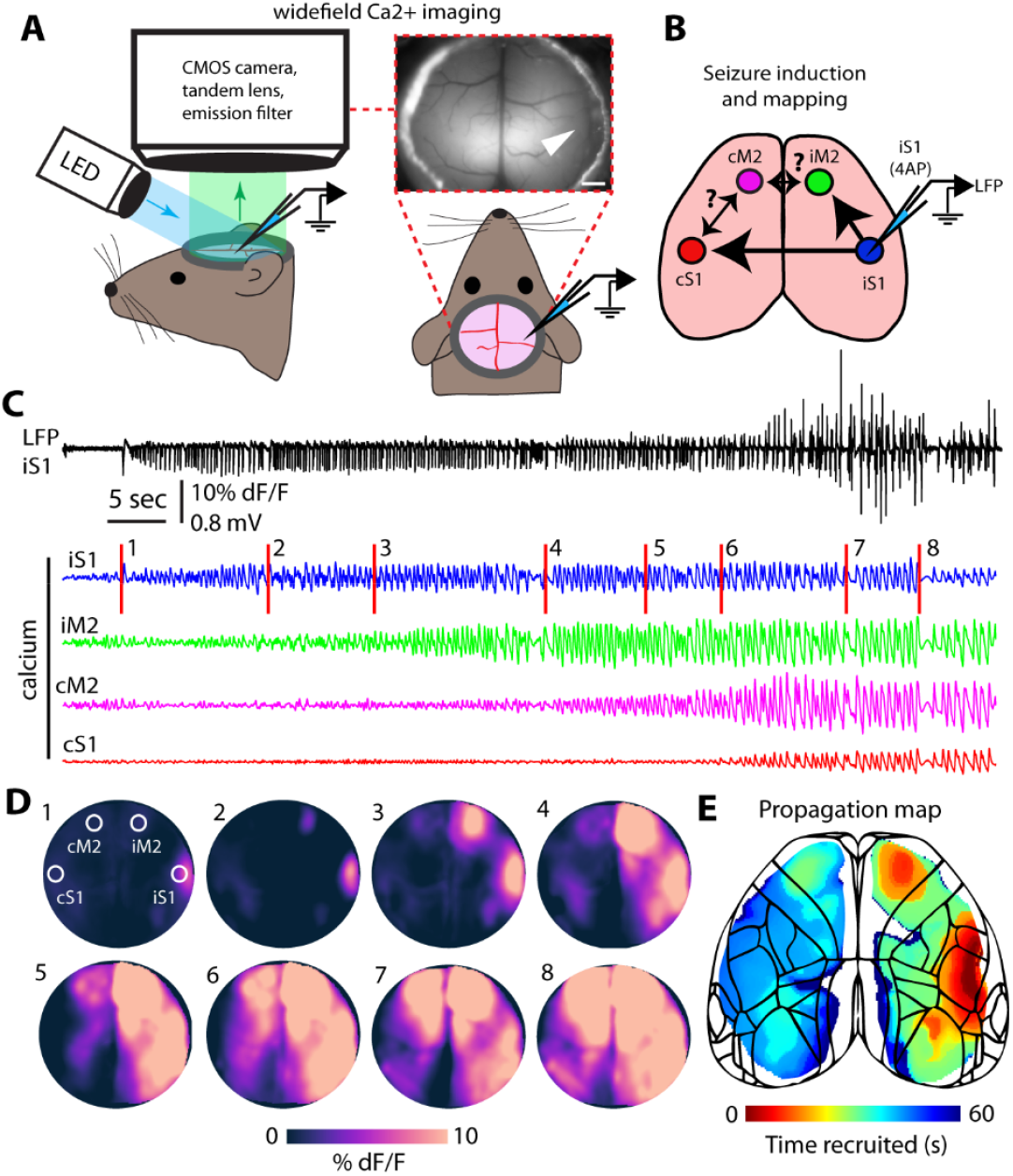
Widefield calcium imaging of the S1 seizure network and its bilateral propagation pattern in dorsal neocortex. A) Transgenic mice expressing the activity dependent fluorophore GCaMP6f in excitatory or inhibitory neurons are implanted with cranial windows and then undergo imaging with simultaneous electrophysiology recording during seizures initiated by 4-AP injection at S1 (white arrow). B) Schematic diagram of the nodes and the possible propagation patterns. C) Electrographic trace (black: LFP) shows an initial spike followed by spike wave discharges. Colored traces (corresponding with colored nodes in B and labelled nodes in D1) show the spread of activity from S1 (calcium fluctuations from ROI in each node). D) Widefield calcium imaging during the seizure, corresponding to red lines in C, show the spatiotemporal propagation of the seizure from iS1 to iM2, to cM2, and finally to cS1. E) Propagation map for this seizure showing a single 2D image of the spatiotemporal spread of the seizure.

In order to differentiate excitatory inhibitory cell involvement in seizure propagation, we imaged GCaMP6f in Thy1+ excitatory neurons (n=9 mice, 4 male) and GCaMP6f in PV+ interneurons (n=7 mice, 3 male). All (n=47) seizure propagation maps were affine-transformed and averaged to create population maps (**Fig. 3A-B**). These heatmaps indicated similar initiation propagation patterns between Thy1+ (excitatory) and PV+ (inhibitory) cell types. Interestingly, as in the single seizure shown in Fig. 2, we observed non-contiguous (saltatory) propagation to iM2. We thus sought to identify whether seizure recruitment across nodes developed by contiguous polysynaptic expansion of the seizure focus, or if activity spread in a saltatory “jump” via monosynaptic long-range network connections^41, 44^. To quantify this, we measured the onset time along a linear track between iS1 and iM2 during seizures to identify the amount of “discontiguity” of the seizure spread (**Fig. 3C**). This analysis revealed that seizures in both Thy1-GCaMP6f and PV-GCaMP6f animals were spreading non-contiguously between iS1 and iM2 (p<.00001 for both Thy1+ and PV+ cells, Wilcoxon signed rank test). In contrast, the propagation discontiguity was significantly lower cM2 to cS1 (p=0.013, Wilcoxon rank sum, n=42 iS1-iM2, n=13 cM2-cS1), suggesting that 4-AP seizures spread more contiguously after crossing the corpus callosum (**Supplemental Fig. 2**).

**Figure 3.**
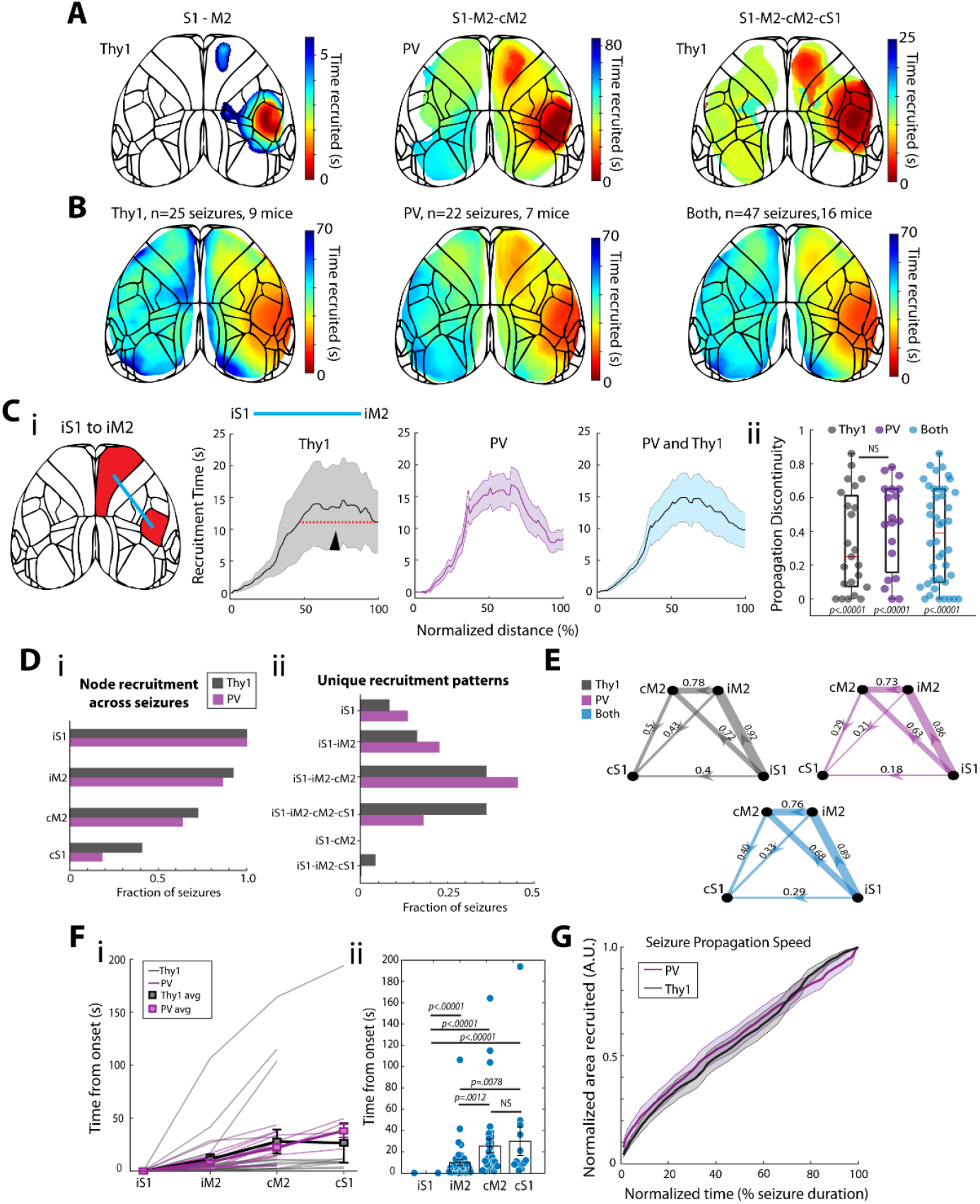
Seizure spread in excitatory and inhibitory cell types. A) Three representative seizure propagation maps show consistent S1 recruitment with variable recruitment of other nodes. B) All seizure propagation maps were affine-transformed and averaged together to create a single “population” seizure map across all seizures. C) Seizures spread outward from S1 but also frequently exhibit non-contiguous propagation to iM2 in both cell types. D) i) Fraction of seizures that recruited each of the major nodes in our study and ii) the fraction of patterns of nodes that were recruited to all seizures. E) Graphs show the “downstream” propagation probability of seizures for Thy1-GCaMP6f and PV-GCaMP6f mice. Arrows indicate “downstream” direction, and the numbers show the fraction of seizures that, if they reach a node, were previously observed in the upstream node. Note that cS1 was the least frequently invaded site, despite direct connection with iS1 (Figure 1). F) i) Both Thy1 and PV activity spread similarly in a stepwise manner across the network. ii) IM2 and both contralateral nodes (cM2 & cS1) were recruited significantly after iS1, although there was no difference between PV and Thy1 recruitment times. G) Propagation speed and area of brain recruited were not significantly different between PV and Thy1. Error bars show SEM.

We next examined the recruitment patterns through the S1-M2 bilateral network (**Fig. 3D-E**). Every seizure initiated in iS1, as expected based on the injection site. IM2 was the most commonly recruited node, followed by cM2 and, lastly, cS1. Notably, when the seizures spread contralaterally, they almost never recruited cS1 without first recruiting cM2 (only one event spread from iS1 to iM2 to cS1, **Fig. 3Dii**). To quantify this trend of node recruitment, we created network graphs of the bilateral S1-M2 nodes and calculated the probability of downstream node recruitment across n=47 seizures (**Fig. 3E**). For both cell types, we observed about 10-15% of seizures remaining focal at S1, and about 90% spreading to iM2. Interestingly, and similar to Figures 1 & 2, we found that when seizures recruited iM2 they had a high probability of crossing the corpus callosum and becoming bilateral “generalized” seizures (78% in Thy1, 73% in PV).

We then compared the temporal propagation patterns between Thy1-GCaMP6f and PV-GCaMP6f mice. As we saw with the propagation maps, seizures initiated at iS1 first spread to iM2 before reaching cM2 and cS1 (**Fig. 3F**). The delay between each node site was significant (p<.00001 for iS1 vs iM2, iS1 vs cM2, iS1 vs cS1; p<.01 for iM2 vs cM2, iM2 vs cS1, Wilcoxon rank-sum test, n=47 S1 seizures, n=42 iM2, n=32 cM2, and n=14 cS1) except for cM2 and cS1 (p>.05) again in agreement with the affine map and discontiguity index data. However, there was no significant difference between Thy1-GCaMP6f and PV-GCaMP6f animals (p>.05 at all node comparisons). We also measured the total brain area recruited across the dorsal cortex during seizures (**Fig. 3G**), which increased over time similarly in Thy1-GCaMP6f and PV-GCaMP6f animals (p=.893, Kolmogrov-Smirnov test, n=25 Thy1, n=22 PV). To investigate the differential role of excitatory and inhibitory cells in ictal onsets, we examined the 5 second period before seizure onset and compared the dF/F at -5 sec and 0 sec relative to seizure onset. An increase in GCaMP signal was observed in both cell types at iS1, but there was no significant difference between Thy1-GCaMP6f and PV-GCaMP6f animals (**Supplemental Fig. 3**, comparing early vs late GCaMP signal: p>.05 for all nodes comparisons; comparing fitted slope during the pre-ictal period: no difference between Thy1-GCaMP6f and PV-GCaMP6f animals; p>.05 for all nodes comparisons, Wilcoxon rank-sum test, n=25 Thy1, n=22 PV).

Our seizure mapping data of the bilateral S1-M2 network highlighted an important role for iM2 in seizure amplification, propagation, and generalization. Thus, we ablated iM2 to see how it might impact contralateral propagation and whether we could redirect the seizure across the sensory component of the corpus callosum (iS1 to cS1). We inserted a metal electrode into right M2 and induced an electrolytic lesion at this site in 4 mice (2 male). This lesion was visually identifiable and altered the S1-M2 network correlations during rest (**Supplemental Fig. 4**). After animal recovery, we then injected 4-AP in iS1 (**Fig. 4**). Our ablation method markedly reduced neuronal activity in iM2 (**Fig. 4B, Supplemental Fig. 4**), and resulted in more contiguous seizure spread (p<.01, Wilcoxon rank sum test, n=42 seizures iM2 intact, n=9 seizures iM2 ablated; **Fig. 4C, Supplemental Fig 4**). Following iM2 ablation, most events (78%) now reached cS1 earlier than cM2 and seizure propagation was altered dramatically. We noted an increase in contiguous spread both in the ipsilateral and the contralateral hemispheres (Supplementary Fig. 4) and a reversal in the propagation in the contralateral hemisphere, with seizures now spreading from cS1 to cM2 and from cM2 to iM2 (**Fig. 4D**). We also noted that, from the affine transformed map, seizure events spread into a medial contralateral region to reach cS1 earlier than cM2 and iM2 (**Fig. 4D-E**). Thus, a primary effect of iM2 ablation was a dramatic alteration in seizure propagation across the network.

**Figure 4.**
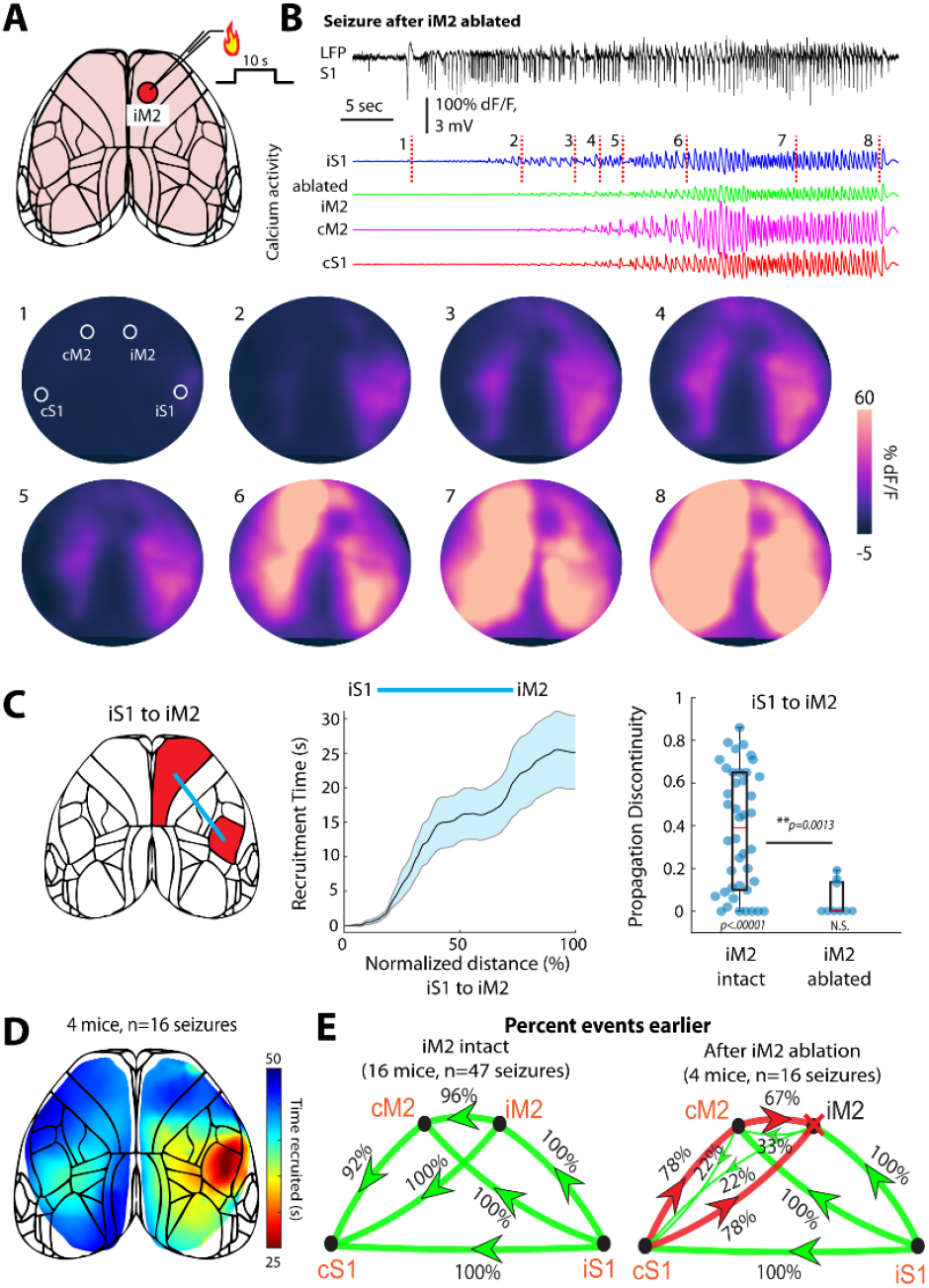
Ablation of iM2 alters bilateral seizure propagation. A) Ablation of iM2 was performed with a lesion induced by 10 s of 1 mA current. B) Seizure activity (LFP and calcium traces) following iM2 ablation by electrolytic lesion and imaging frames (below: numbered frames correlate with the red dashes). Note that some fluorescence still occurs at iM2 but is notably reduced following ablation (also see Supplemental Fig. 4). C) Following iM2 ablation, seizures spread with more contiguity between nodes. D) Affine-transformed average propagation map shows how seizures spread following iM2 ablation. cS1 recruitment tends to be earlier than iM2, and cM2, but the seizures reach cS1 through an indirect route. E) Before ablation (16 mice, 47 seizures), seizures spread with high rates from iS1 to iM2 to cM2 to cS1, but these patterns changed following iM2 ablation. Red lines in ablation graph (right) show where a change in seizure direction occurred following iM2 ablation (4 mice, n=16 seizures).

We further investigated the inhomogeneity in cross-callosal seizure propagation (i.e. preferential spread anteriorly between iM2 and cM2, rather than posteriorly between iS1 and cS1) and whether this was a phenomenon unique to seizure activity by using direct sub-ictal electrical stimulation of iS1 (**Fig. 5A-B**). Stimulus durations were set to loosely simulate the duration of interictal events (100 ms), polyspike events (1 sec), and ictal events (10 secs) to determine if duration was a factor in excitatory-inhibitory cell recruitment. Each stimulus parameter produced reliable calcium imaging responses across the S1-M2 network in dorsal cortex (n=10 animals, 5 Thy1-GCaMP6f, 3 male, and 5 PV-GCaMP6f, 3 male). We observed a notable difference in how cM2 and cS1 regions responded during stimulation in the two cell types. Stimulation of Thy1-GCaMP6f animals generally resulted in a strong response at cM2 and relatively weaker response at cS1; meanwhile, PV-GCaMP6f activity was relatively stronger in cS1 than cM2 (see seed correlation maps in Fig. 5A-iv). We quantified this effect over the 3 stimulation durations by plotting the cS1 and cM2 responses for each trial (**Fig. 5C**), finding that the cS1 bias of PV-GCaMP6f and the cM2 bias of Thy1-GCaMP6f signals was significant at all durations (100 ms, cM2 vs cS1 p<.00001; 1 sec, p<.00001, 10 sec, p<.00001, Chi-square proportion test; cM2/cS1 ratios, comparing PV vs Thy1: 100 ms, p<.01; 1 sec, p<.00001; 10 sec, p<.05,

**Figure 5.**
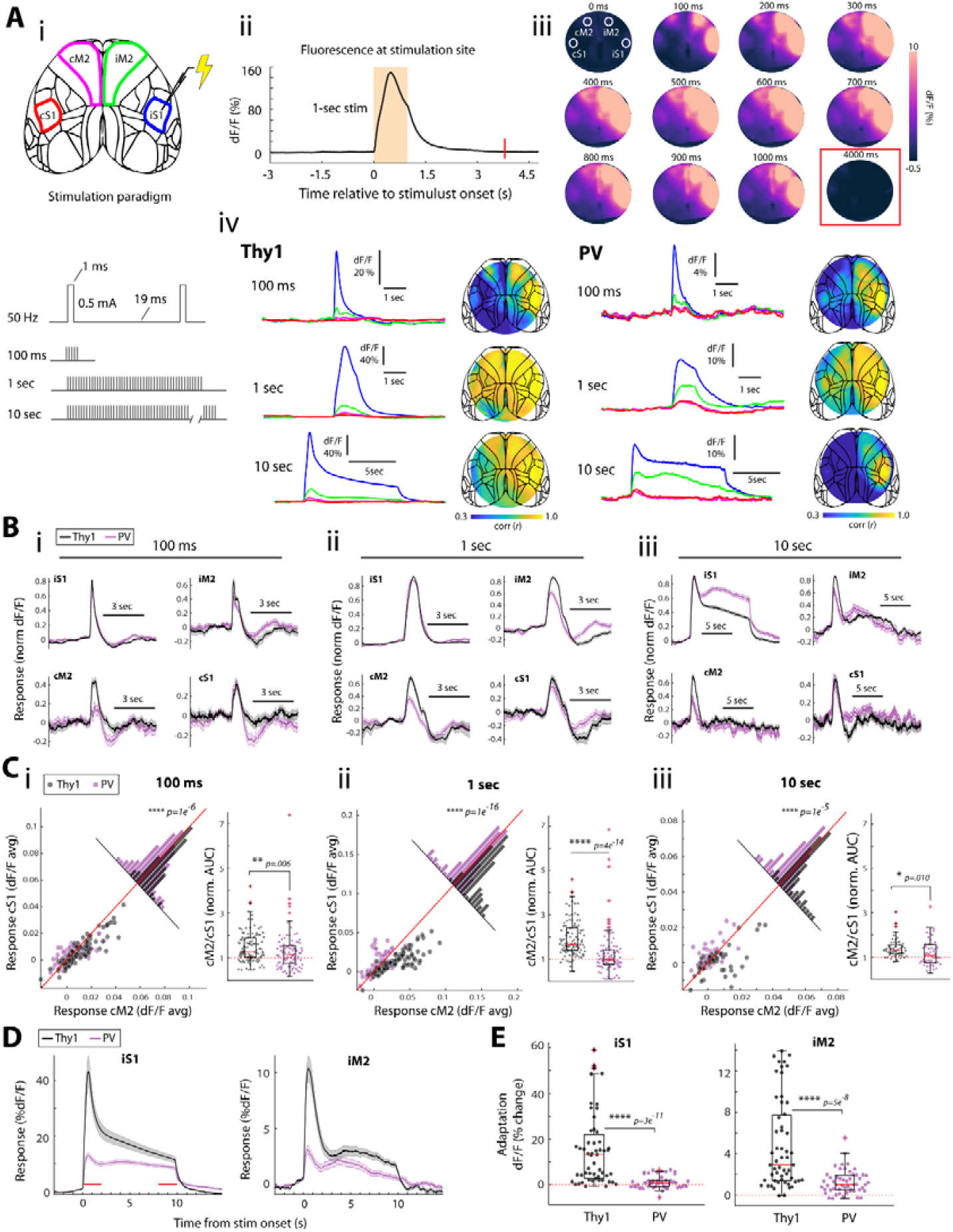
Focal electrical stimulation recapitulates seizure propagation patterns and identifies differences in Thy1 and PV cell recruitment. A) i) Electrical stimulation trains (0.5 mA, 50 Hz) were applied at S1 for three different durations. ii) GCaMP6f activity was recruited rapidly. iii) Imaging frames show the S1-M2 network recruitment during 1-sec stimulation, and a return to baseline within 4 seconds of stimulation termination. iv) Example stimulations of 100 ms, 1 sec, and 10 sec in Thy1-GCaMP6f and PV-GCaMP6f mice. Maps show S1-seeded correlations for all pixels in the imaging window averaged over all stimulation trials: Thy1 n=22, 21, and 10 trials for seed maps 100 ms, 1 sec, and 10 sec; PV n= 21, 23, and 10. B) Average and standard error (shading) of GCaMP6f activity at the node sites for all stimulations in 8 mice (n=4 Thy1-GCaMP and n=4 PV-GCaMP animals, 100 ms, 1 sec, and 10 sec trials n=109, 107, and 55 for Thy1, n=101, 110, 52 for PV). C) GCaMP response at cS1 and cM2 following 100 ms (i), 1 sec (ii), and 10 sec (iii) stimulations. Values above the line of unity indicate stronger response in cS1 compared with cM2. The Thy1 vs PV difference was significant across stimulation durations. Boxplots show cM2/cS1 response ratios calculated on the normalized area under response curves. D) Calcium response from 10-sec stimulation trials are shown, highlighting a difference in the sustained activity of PV-GCaMP6f, while Thy1-GCaMP6f activity tended to decay, or adapt, over time (this can also be seen in B-iii). Red lines indicate periods of early and late activity to calculate adaptation. E) Adaptation was significantly higher in Thy1-GCaMP6f vs PV-GCaMP6f at iS1 and iM2 (n=55 Thy1, n=52 PV trials).

Wilcoxon rank-sum test. Non-normalized response data are shown in **Supplemental Fig. 5A**). In agreement with this, we also observed significantly higher correlation between response at iS1 and cM2 for Thy1-GCaMP compared to PV-GCaMP (p<.0001 for all stimulation durations, Wilcoxon rank-sum; **Supplemental Fig. 5B**). At the longest duration, 10 sec, we also observed a difference in the response profile (shape of calcium trace) between Thy1-GCaMP6f and PV-GCaMP6f animals. At S1, Thy1-GCaMP6f responses generally showed an initial strong response followed by a slow adaptation or decay, while PV-GCaMP6f responses showed an initial response followed by a comparatively well-maintained responsiveness throughout the 10-sec stimulus train (**Fig. 5D**). To measure this between Thy1-and PV-GCaMp6f animals, we calculated the relative adaptation during the 10 sec stimulation in iS1 and iM2, finding a significantly higher adaptation in Thy1-GCaMP6f compared with PV-GCaMP6f responses (**Fig. 5E**, iS1, p<.00001; iM2, p<.00001, Wilcoxon rank-sum test).

These stimulation experiments highlight the functional connectivity of the S1-M2 network and suggest that E/I balance may bias seizure spread from iS1 to iM2 and away from cS1. However, these stimulations were brief, and do not accurately recapitulate the dominant frequency of ictal events. Further, the 10-second stimulations revealed that Thy1 activity was more likely to adapt to a 50 Hz stimulus over time while PV inhibitory activity was relatively robust throughout this period. We thus hypothesized that 50 Hz recruitment of PV interneurons could lead to a slow adaptation of local excitatory neurons through constant inhibitory neurotransmitter release. In seizures, this process has been proposed to produce a change of the chloride reversal potential in excitatory neurons, turning GABA into a (locally) excitatory neurotransmitter^45-47^. To test this possibility and better understand how seizure-like discharges affect a bilateral network, we developed a “seizure-like” electrical stimulation pattern to apply during widefield imaging.

We first used our 4-AP seizure LFP data to identify the frequency with strongest power, which, across all seizures, was 2.88 Hz (**Fig. 6A; Supplemental Fig. 6**). Based on this, we created an electrical stimulation pattern that provided a 3 Hz seizure-like stimulation, which robustly activated S1-M2 network activity across the dorsal cortex (**Fig. 6B**). We applied this seizure-like stimulation for 10-second periods in 6 mice (n=3 Thy1-GCaMP6f, 1 male, and n=3 PV-GCaMP6f, 2 male) during simultaneous widefield calcium imaging. Unlike the continuous 50 Hz stimulation (Fig. 5), for both cell types, we observed a facilitating response to the 3 Hz stimulation at iS1 and iM2 (**Fig. 6C-D**; adaptation values for iS1: Thy1, p<.05, PV p<.01; for iM2, Thy1 p<.05, PV p<.01; Wilcoxon rank-sum test; n=12 Thy1 stimulations, n=11 PV; contralateral node data can be seen in Supplemental Fig. 6). We also examined whether any mismatch of cM2 and cS1 activity was present, but, unlike the continuous 50 Hz stimulation, we did not observe any significant bias of activity for cM2 vs cS1 (**Supplemental Fig. 6D**). We did, however, observe a significantly stronger correlation between iS1 and iM2 vs cM2 for Thy1-GCaMP6f (p<.01 Wilcoxon rank sum; **Fig. 6E**) and a significantly stronger correlation between PV-GCaMP6f responses at cS1 vs Thy1-GCaMP6f responses, suggestive of a relatively higher inhibitory recruitment at cS1 relative to cM2 (an effect like we observed in our 50 Hz stimulation, Fig. 5C). We also extended our stimulations to 30-second durations (**Fig. 6F**). This resulted in a similar pattern of results as with 10 second stimulations, showing significant facilitating response at iS1 and iM2 but with no difference in PV-GCaMP6f vs Thy1-GCaMP6f (**Fig. 6G;** Thy1: iS1 p<.05; iM2 p<.01; PV: iS1 p<.01, iM2 p<.01; No significant difference between Thy1-PV, Wilcoxon rank-sum test, n=13 Thy1, n=11 PV trials). We also observed, as with the 10 sec stimulation, a significantly stronger correlation between iS1 and cM2 versus cS1 for Thy1-GCaMP (**Fig. 6H**, p<0.05, Wilcoxon rank sum), though no significant differences between Thy1 and PV responses. Thus, the 3 Hz seizure-like stimulations did not evoke a difference in adaptation between Thy1 and PV neuronal responses as we saw with 50 Hz continuous stimulation. Rather, we observed that these seizure-like frequencies evoked robust facilitating activity in both cell types.

**Figure 6.**
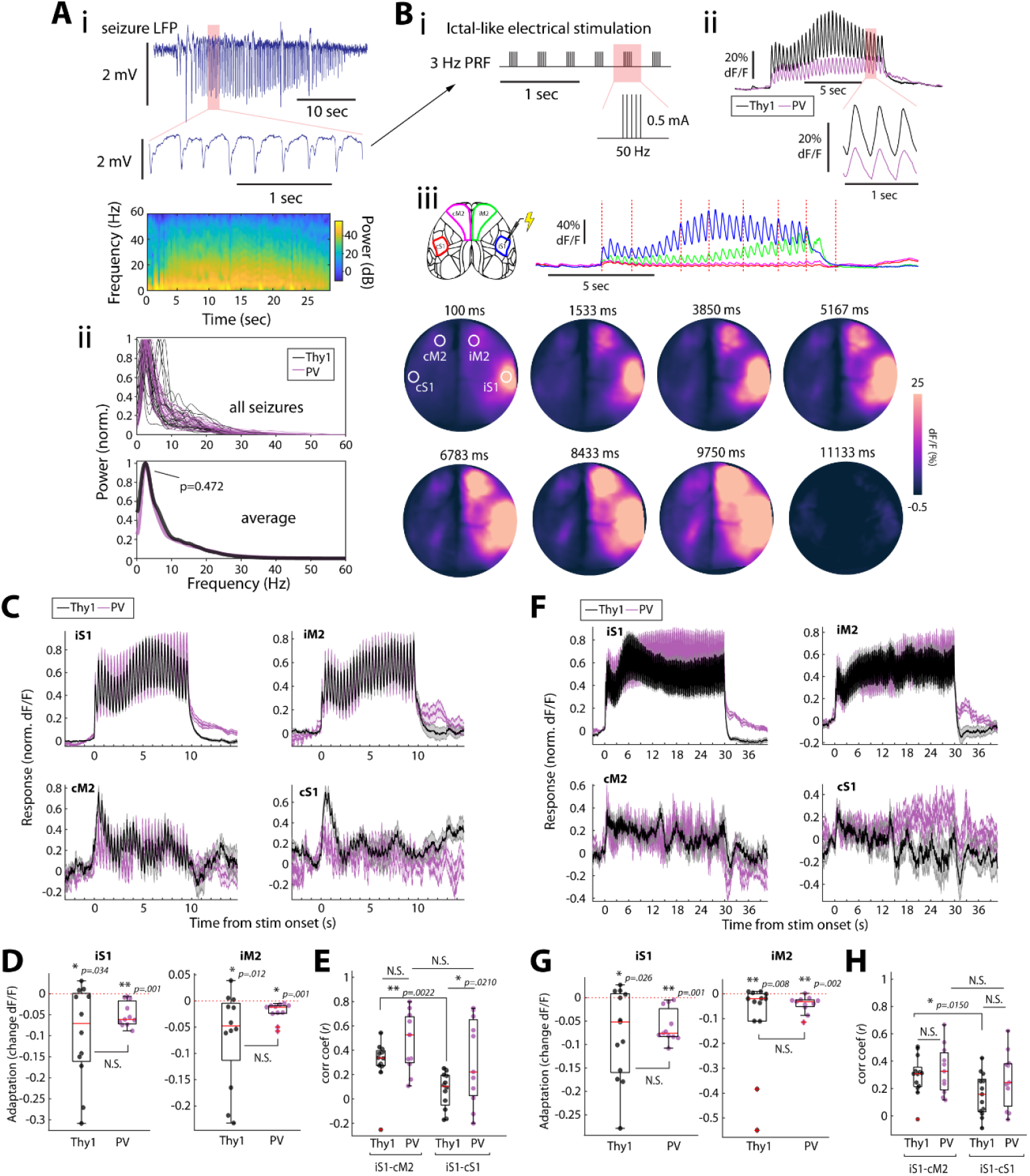
Seizure-emulating electrical stimulation recruits network sites. A) i) Sample seizure data showing low frequency spike-wave discharges. ii) Power spectra from electrographic data for all seizures are shown, with a peak frequency of 2.88 Hz, which was not significantly different in Thy1-GCaMP6f and PV-GCaMP6f animals. B) i) Seizure-like stimulation was created by using a 3 Hz pulse response frequency with 100 ms 50 Hz bursts. ii) Bursts reliably recruit neural activity at the stimulation site. iii) Widefield imaging shows immediate S1 recruitment and early iM2 recruitment during 3 Hz ictal-like stimulation, which spreads to contralateral sites during this 10-second stimulation. C) Averaged normalized responses to ictal-like stimulation show a facilitating response in iS1 and iM2, with weaker responses in contralateral cortex. D) Both Thy1-GCaMP6f and PV-GCaMP6f exhibit significantly facilitating responses throughout the stimulation at iS1 and iM2 but were not significantly different from each other. E) The correlation of iS1 and cM2 was significantly higher for Thy1 responses compared to iS1 and cS1, while PV responses were significantly more correlated between iS1 and cS1 compared with Thy1 responses. F) Same as C but for 30 second ictal-like 3 Hz stimulation trials. G) As with 10 second stimulations, the 30-second stimulation did not differ significantly between Thy1-GCaMP6f and PV-GCaMp6f mice, but both showed significantly facilitating responses. H) As with 10 second stimulations, 30-sec stimulation showed a higher correlation between iS1 and cM2 vs cS1 in Thy1 (but not PV) activity.

Neurostimulation experiments at 50 Hz highlighted a difference in E/I, but at longer duration also showed a relatively stronger adaptation of Thy1 excitatory activity. When we turned to 3 Hz stimulation this effect disappeared, and both PV and Thy1 cell types exhibited robust facilitation. Based on this finding, we returned to our seizure data and calculated the relative adaptation in Thy1 and PV experiments and examined this alongside the stimulation data (**Fig. 7A;** n=9 Thy1 mice, n=7 PV mice; stimulation data averaged within mice before affine-transforming, n=5 Thy1 and n=5 PV mice for 50 Hz; n=3 Thy1 and n=3 PV mice for 3 Hz; n=9 Thy1 and n=7 PV mice for seizure data). Seizures, like our 3 Hz data, exhibit robust facilitation (measured in the earliest 20% vs last 20% of seizure, see Methods). This effect was seen across nodes during seizures (**Fig. 7A-B**), confirming a link between low frequency activity initiated at a focal site and subsequent brain-wide facilitation of both excitatory and inhibitory cell types.

**Figure 7.**
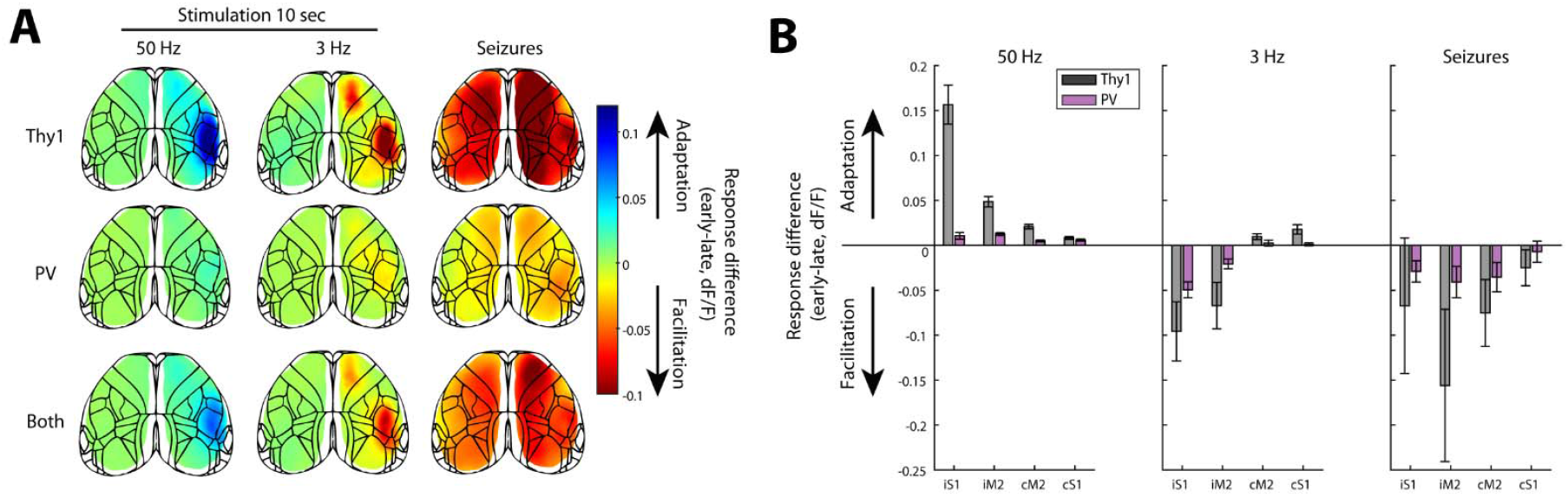
Adaptation and facilitation during stimulation and seizures. A) Average changes in early vs late response to 50 Hz or 3 Hz stimulation, and seizures, across cell types. B) Average response differences shown at S1-M2 bilateral node sites. Error bars show SEM.

## Discussion

By inducing focal seizures and imaging neural activity within a well-described multi-nodal, monosynaptically-connected bi-hemispheric network, we show that seizures spread inhomogeneously, in both a non-contiguous saltatory and contiguous Jacksonian manner, along preferential pathways based on E/I-specific cell recruitment. In the S1-M2 network, seizures starting in S1 preferentially spread anteriorly to M2 and across the corpus callosum in motor regions, rather than to the contralateral S1 mirror sensory region. Thus, the frontal M2 area of the brain and its callosal connections appear to act as an ictal amplifier for seizure spread and generalization compared with the somatosensory areas. With focal stimulation of S1, we show that neuronal activation in S1 alters E/I balance in M2 in favor of excitation but favors inhibition in contralateral S1. Hence, inhomogeneous long-range recruitment of specific cell-types, or “functional” connectivity may explain ictal propagation pathways over mere “anatomic” connectivity.

In the bilateral S1-M2 network, the M2 node functioned as a seizure amplifier. Ablation of M2 radically altered seizure propagation. Nevertheless, when iM2 was ablated, seizures still spread contralaterally although they did not follow the most direct anatomic connection across the corpus callosum to S1. Rather, they spread to adjacent areas that eventually spilled over to S1. This was unexpected given the connectivity of S1 with contralateral S1. We thus applied electrical stimulation to S1, mapping excitatory (Thy1-GCaMP6f) and inhibitory (PV-GCaMP6f) activity across the network. We found that stimulation robustly recruited the bilateral S1-M2 network, but with markedly different functional recruitment: E/I balance was higher at the frontal cM2 (favoring excitation) node compared with cS1 (favoring inhibition). This functional excitatory connectivity may explain why seizures consistently spread across the frontal iM2-cM2 route. When we examined a longer stimulation duration (10 sec) we also observed a significant difference in PV-GCaMP6f and Thy1-GCaMP6f responses over time at ipsilateral network sites, suggesting that longer periods of activation can differentially affect E and I activity. This led us to next investigate whether this response difference could occur during seizure-like stimulations as well. We thus identified the dominant discharge frequency of our seizures and created a “seizure-like” 3 Hz stimulation pattern. Contrary to expectations, we did not observe adapting excitatory responses during this stimulation. Rather, both PV-GCaMP6f and Thy1-GCaMP6f responses exhibited facilitation throughout the stimulation period, even at a downstream ipsilateral node site. These findings reveal a significant difference in the effects of high versus low frequency stimulation, which can differentially elicit excitatory or inhibitory responses.

### Neurostimulation seizure therapy

Electrical neurostimulation can be effective at reducing seizure frequency while conferring less damage to neural tissue compared with resection, which can mitigate morbidity and potential side effects^48^. Yet, complete seizure freedom is rarely achieved^49^. Nevertheless, advancements in these tools, both conceptually and methodologically, could improve patient outcomes^18^. Despite extensive modelling^50, 51^, the network effects of focal stimulation have not been examined experimentally. Our findings demonstrated that different activation frequencies can differentially recruit excitatory and inhibitory activity across a cortical network. Our cell-type-specific imaging data suggests this desynchronization may rely on differential response to high frequency stimulation such that PV interneurons, which encompass a major class of fast-spiking interneurons^52^, are more robustly recruited to fast current changes while excitatory pyramidal cells exhibit a slowly adapting response. Evidence exists for this in a recent study by Hughes et al.^40^ in which pyramidal cells were significantly more likely to show a slow decay, or adaptation, throughout prolonged intracortical stimulations (up to 100 Hz). Those authors posited that recruitment of inhibitory cells during stimulation could be the mechanism by which pyramidal cell responses show adaptation: as inhibitory GABA is released repeatedly, local excitatory cells will become less responsive over time. Our data show that low frequency stimulation of 3 Hz led to a robust recruitment of both cell types. Most importantly, these effects propagate, presumably along direct axonal pathways, to downstream network sites (though diffuse effects are also possible^53^, particularly through common nodes within a network^54^).

### Connectome vs. observed seizure propagation

Some recent studies have suggested that personalized neurostimulator planning may be optimized for individual patients with drug-resistant epilepsy^5, 11, 12, 55, 56^. In this vision, a patient would undergo imaging and sEEG to thoroughly identify the network features involved in his or her seizures. From this data, a personal network model would be created and then the ideal stimulator site and parameters could be identified based on testing how the model responds to different stimulation locations^57^. Towards this goal, a recent study created a “virtual twin” of 30 epilepsy patients, combining anatomical and functional data from these individuals to create models that permitted comparison of simulated and observed seizure data, which revealed impressive similarities that could potentially guide surgical decisions^50^. Our findings here present a potential caveat: while the structural and functional architecture of a brain network is visible at rest, a seizure may propagate through sites that are difficult to identify based on this data. Our simple cortical network involved only four nodes, and we hypothesized seizure propagation would occur by one of two routes. However, we observed faster seizure spread to contralateral cortex by an unexpected route, and it was only during electrical stimulation with cell-type-specific imaging that we could identify possible mechanisms for this propagation pattern. Thus, relying on connectome (structural and resting state) information may be insufficient to produce a fully comprehensive “virtual brain” model that identifies seizure propagation routes. While application of focal stimulation, as was included in the recent virtual twin study^50^, could improve identification of seizure propagation patterns, predicting the optimal neurostimulator parameters and location may require better understanding of how E/I balance is affected across the brain by focal neurostimulation.

### Limitations

One limitation of this seizure propagation mapping is that we have no deep brain imaging. Our widefield mesoscale imaging is fast and encompasses broadly distributed bilateral areas, but the existence of other seizure nodes, such as the thalamus and hippocampus, cannot be confirmed with our current methods. Indeed, the S1-M2 network is directly connected with the ventral posteromedial thalamic nucleus (VPM). Our seizure propagation maps, particularly after ablation of iM2, revealed diffuse seizure recruitment in contralateral cortex, and we speculate that this occurred through deep brain structures (e.g., the VPM or the hippocampal commissure). Another limitation of our imaging is that the E/I imbalance we observe is limited to two populations of genetically defined cells: Thy1+ and PV+ neurons. While both cell types have been described as encompassing multiple classes of either excitatory or inhibitory cells, these are still only subpopulations. For example, multiple subgroups of PV+ interneurons have been well-characterized: chandelier and basket cells show differential preference for axon initial segment-and somatodendritic-targeting of excitatory neurons, respectively^52, 58, 59^. Though PV+ interneurons are expected to provide the majority of inhibitory tone onto excitatory neurons, it is certain that other interneurons, such as somatostatin+ cells^60^, also contribute to this inhibition. Our use of 4-AP is also a limiting factor: while this chemical induces robust focal seizures, it does not perfectly emulate the seizure initiations that would occur in human patients. Finally, our study may be limited in its relevance to other brain networks. We chose the S1-M2 bilateral network for its well-characterized nature and accessibility for mesoscale investigation, however, it is likely that other networks, particularly subcortical ones^25, 26^, may exhibit other features that we cannot observe in the neocortex.

## Materials and Methods

Experimental procedures were approved by the Weill Cornell Medicine Institutional Animal Care and Use Committee following NIH guidelines.

### Animals and surgeries

Data was collected in mice (6-8 weeks age) expressing GCaMP6f in Thy1-positive pyramidal neurons (C57BL/6J-Tg(Thy1-GCaMP6f)GP5.5Dkim/J, Jax 024276) or parvalbumin-positive (PV) inhibitory neurons (B6.129P2-Pvalb^tm1(cre)Arbr^/J, Jax 017320, homozygous, crossed with GCaMP6f reporter B6J.Cg-Gt(ROSA)26Sor^tm95.1(CAG-GCaMP6f)Hze/^MwarJ, Jax 028865). These mice are hereafter referred to as “Thy1-GCaMP6f” and “PV-GCaMP6f”, respectively. Animals (of both sex) aged 6-10 weeks were implanted with cranial windows: mice were first anesthetized with 3.5% isoflurane mixed with pumped ambient air and injected with 2 mg/kg Meloxicam. Bupivacaine (0.1 mL,0.25%) was locally injected under the scalp. Body temperature was maintained at 37 degrees Celsius with an electric heating pad (Harvard Apparatus, USA). Next, the animal was head-fixed in a stereotactic frame (Kopf, USA) and placed in a nose cone that delivered 1-1.5% isoflurane throughout the surgery. Hair was removed by scissors and an 8×9 mm skin opening was made. Next, a craniectomy was performed over the skull using a dental drill and size 005 bur. A circular window of 8.2 mm diameter was then implanted over the exposed cortex and attached at the edges with Vetbond (3M, USA). Next, a metal head plate (Neurotar, FI) was implanted over this window and adhered to the skull with dental cement (C&B Metabond, Parkell, USA). Mice were provided 2 mg/kg meloxicam daily for 3 days following surgery and monitored for any adverse health conditions.

### Seizure induction

We leveraged the known white matter connections revealed by the Allen Atlas map **(Fig. 1A)** to create a multimode neocortical seizure network. Following animal recovery of one week, seizures were induced by injection of 1 mM 4-Aminopyridine (4-AP) into right S1 (-1.7 AP, -3.5 ML), at a depth of 300 microns from the brain surface (targeting layer 2/3) (**Fig. 1B**). This was performed with a glass micropipette electrode, injecting 4-AP in 10-nL quantities (∼100 nL) until focal seizures were observed on the simultaneously colocalized electrographic recording.

### Imaging

Mice were head-fixed in an apparatus that permitted free running (Neurotar, Finland). Widefield calcium imaging was performed with a CMOS camera (xiC, Ximea, USA; with tandem lens) running at 120 Hz, with frames capturing fluorescence following illumination by 470 and 530 nm via light-emitting diodes and optic fibers (Thorlabs, NJ, USA) at a functional imaging rate of 60 Hz for GCaMP6f. Further details can be found in Yang et al^61^.

### Electrophysiology recordings

Electrophysiological data was recorded from each of the four nodes in our bilateral cortical network from glass micropipette electrodes (**Fig. 1B;** ∼1 MΩ). Local field potential (LFP) was recorded at 1 kHz sampling with a 1-500 Hz filter. The electrophysiology system consisted of a preamplifier (headstage, x1000) and second-stage amplifier that included bandpass filtering 1-500 Hz (A-M Systems 1800, USA) and a digitizer (CED 1401, Cambridge, UK) before arriving at a PC running Spike2 (CED, UK). The LFP was recorded at 1 kHz and analyzed offline (described below). Electrographic data was stored locally on a PC until post-experiment processing. Files were opened and analyzed in MATLAB (v2021a, The Mathworks, USA). These signals were recorded in files that also recorded TTL pulses during microstimulation trials (a TTL pulse was recorded for every electrical stimulation pulse) and frame times from our imaging camera to allow alignment with GCaMP data below. For spectral power calculations, we used MATLAB’s *pspectrum()* (using Welch’s method for power spectrum estimation). This same function was used for power calculation in calcium data described below, with frequency limits set to values seen in figures (e.g., 0 to 30 Hz, Supplemental Fig. 6; LFP was recorded at 1 kHz frequency while calcium signals were all recorded at 60 Hz).

### Cortical ablation

Electrolytic lesions were induced in neocortical M2 following parameters described by other studies^62, 63^. After anesthetizing the mouse, an electrode was inserted through the PDMS film window into the neocortex. This electrode was epoxy-coated such that 0.5 mm of tip was exposed to brain tissue. Using a Master-8 and ISO-flex system (AMPI, IL), a constant current of 1 mA (anodal) was applied for 10 seconds. These parameters create an immediate visible 1 mm diameter discoloration of the brain at the stimulation site. Animals were given meloxicam (2 mg/kg) after the procedure and for 3 days post-lesion. No obvious symptoms or negative health issues were noted in any of the animals.

### Microstimulation

Stimulation experiments were performed with a glass micropipette electrode with a ∼50 µm tip opening. The pipette was filled with 0.9% sterile saline and placed over a stainless-steel wire. Next, the electrode was inserted into S1 (depth 300 microns). And attached to the ISO-flex and Master-8 electrical stimulator system. Unlike the ablation experiments, here we used a 0.5 mA anodal stimulation pattern (varying the duration and frequency as described in Results). During 50 Hz stimulation, a 1-ms pulse was followed by a 19 ms rest period (trials were randomly varied at either 100 ms, 1 sec, or 10 sec). 3 Hz stimulation was achieved with this same pattern but treating each 5-pulse (100 ms) period as a single pulse with a pulse repetition frequency of 3 Hz (10 and 30 sec randomly varied trials). Stimulations were delivered and TTL pulse carried a corollary signal to the Spike2 PC for offline identification. All stimulation trials were randomized and provided with 15-30 seconds between trials, a duration that always showed a return to baseline signal in our calcium imaging data.

### Calcium imaging data analysis

Data processing and statistics were performed with MATLAB 2021a (The Mathworks, USA). Calcium imaging data frames were deinterleaved from the 530 nm frames. Next, to counteract changes in GCaMP caused by hemodynamic contamination, a modified Beer-Lambert equation with a path length correction factor was applied to the imaging data using the equation:

Where *Ftrue* is the signal associated with GCaMP after removing hemodynamic bias at timepoint *t*, and *F* is the observed fluorescence, *tb* is the baseline, and *I* is the intrinsic optical signal from 530 nm illumination (after application of a 2 Hz lowpass Butterworth filter). Following this hemodynamic correction, we calculated the change in GCaMP fluorescence within individual seizures and stimulation trials using the equation:

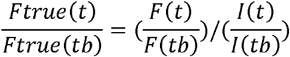

Where *Ft* is the hemodynamic-corrected activity at frame *t* and *Fb* is the baseline fluorescence. When comparing dF/F, traces were smoothed by using a moving average of 50 ms. Area under curve of the calcium signal was calculated using the *trapz*() function in MATLAB.

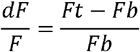

### Seizure propagation

After GCaMP was hemodynamic-corrected and converted to dF/F, we extracted this calcium change at individual brain network sites based on anatomic locations of our network nodes. S1 was centered at -1.7 posterior, +3.5 lateral, while M2 was centered at +1.0 anterior and +0.75 lateral (all coordinates are relative to Bregma and in units of mm). A 3×3 pixel region was centered around these centroids and averaged per frame to extract GCaMP signal from our network nodes. For mapping of seizure activity, we applied our analyses to all pixels in the field of view (FOV) in the PDMS film window. We first compared seizure onset detected in calcium signals with onset detected in the electrographic recording to identify the optimal 2D mapping method (Supplemental Figure 1). We compared “line length” (LL), theta band (5-8 Hz), delta band (1-4 Hz), and root mean square (RMS) methods. We first applied a high pass filter (*highpass()*, 1 Hz cutoff) in MATLAB. Then for each calculation, we identified the average and baseline signal (e.g., average and baseline LL) in a 3 second period before electrographic seizure onset. We then identified, for each individual pixel, when the signal crossed 2x this value, which we used as the threshold for comparing each method (see supplemental fig 1 for data from this comparison across animals and seizures). We found that line length was a robust detector of calcium signal associated with an electrographic seizure and thus used this method to calculate propagation maps. These maps were simply the times at which each pixel was recruited to the seizure; pixels that did not cross the threshold were considered “not recruited”. To compare Thy1 and PV seizure propagation speed, we calculated the normalized area and normalized time of recruitment for each individual seizure, converting each to percentile.

To calculate the contiguity of seizure spread, we created a “discontiguity index” by identifying all pixels in a straight line between iS1 and iM2 and cS1 and cM2 (see Supplemental Fig. 2). This line was made with a MATLAB implementation of Bresenham’s line algorithm^64^ and then, for each animal, the trace of recruitment time was identified for each of these pixels and then normalized to percentile for comparison across seizures. These traces could then be averaged together to quantify the discontiguity during propagation over the FOV between selected network sites.

Affine transformation was performed by first applying a 2D gaussian filter (*imgaussfilt()* in MATLAB) with a sigma of 1 pixel to all propagation maps. Next, images were registered to a starting map by aligning each of the 4 network nodes (iS1, iM2, cM2, cS1) and then applying *fitgeotrans()* followed by *imwarp()* with *imref2d()* in MATLAB to align both of the propagation maps. This was done for every seizure propagation (aligning to the same registering map) and then these maps were averaged together to create single population propagation maps. Seed correlation was calculated as Pearson’s correlation coefficient *(r)*, and seeding was performed by selecting the dF/F at one network site (e.g., as indicated in Supplemental Fig. 4) and then calculating the correlation coefficient with every other pixel in the FOV.

### Stimulation data

Microstimulation trials with no observed current in the adjacent electrographic recording and trials with dF/F change at the stimulation site <3x the baseline calcium activity were omitted (9 of 543 trials were omitted for 50 Hz stimulation, and no trials were omitted for 3 Hz stimulation trials). The degree of adaptation and facilitation of neural responses following 10-and 30-second stimulation trials was calculated with a method adapted from Hughes et al.^40^: we identified the initial and final response during stimulation trials, calculating early minus late periods (2 sec) to identify the amount of adaptation as change in dF/F over time. This metric was calculated on 10-second stimulation at 50 Hz (continuous) as well as the 3 Hz “seizure-like” data. For seizure data we used the earliest 20% and last 20% of seizure period for this calculation.

## Acknowledgments

Supported by the Citizens United for Research in Epilepsy (CURE Epilepsy no. CURE241473, JEN) and the Mitchel Alan Ross Grant for Epilepsy and Asphyxia research from Weill Cornell Medicine (JEN). The authors also thank the veterinary and animal care staff at Weill Cornell Medicine and members of the Department of Neurosurgery for their assistance and discussions of this manuscript.

