## Supplemental data for "Functional rather than anatomic connectivity predicts seizure propagation in a multi-node model of focal neocortical epilepsy"

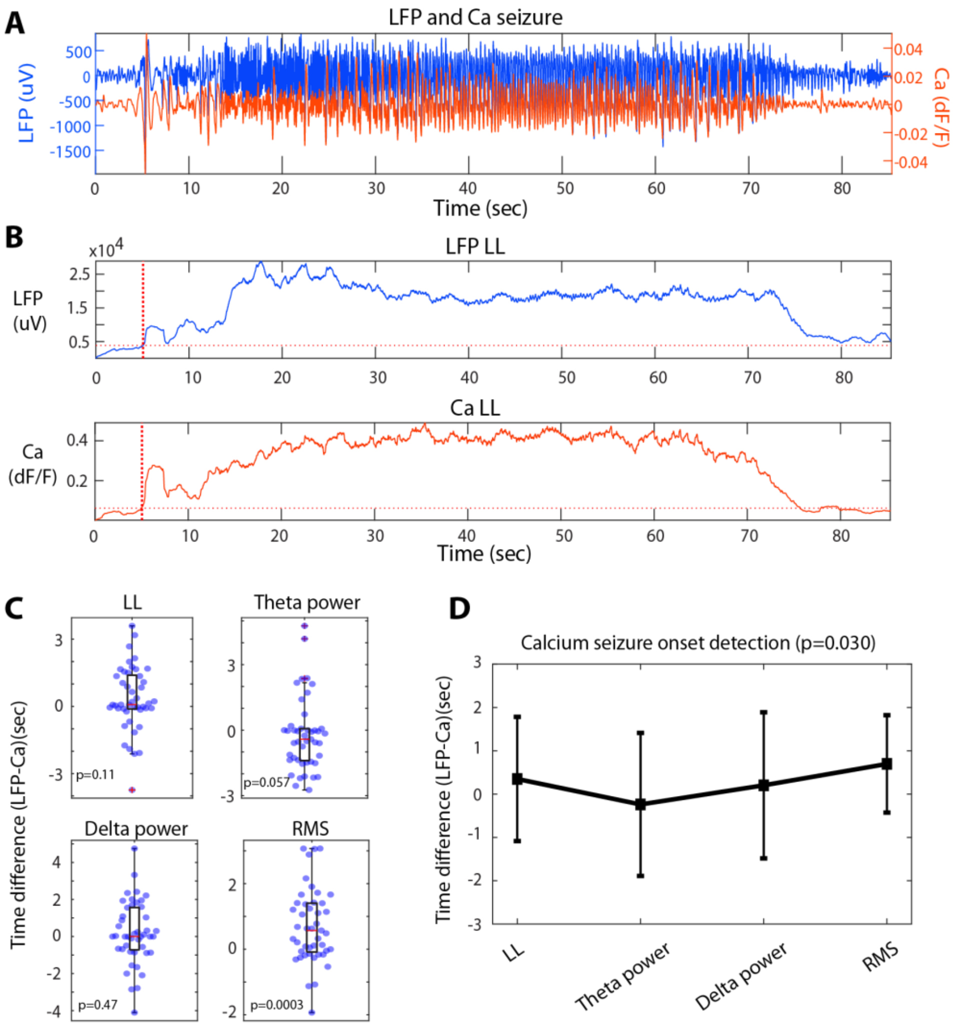


**Supplemental Figure 1.** Pixel-wise seizure onset detection. A) LFP and GCaMP (Ca) activity at S1 during a seizure (high pass filtered >1 Hz). B) Line length is calculated in a moving 2-sec window, showing general agreement between LFP and Ca traces (blue and orange, respectively). Onset was identified from significant deviation from the baseline period. C) We tested line length (LL) mapping and tested theta power (1-8 Hz), delta power (1-4 Hz), and root mean square (RMS), comparing LFP and Ca onset detection. RMS was the only method significantly different from zero (p=.00003, Wilcoxon sign-rank test, n=45 seizures). D) Comparing LL, Theta, and Delta: Delta and Theta were closer to zero, but were not significantly different from LL. Given that LL showed a slightly lower variance, and was computationally faster, we chose to use this method to create seizure propagation maps.

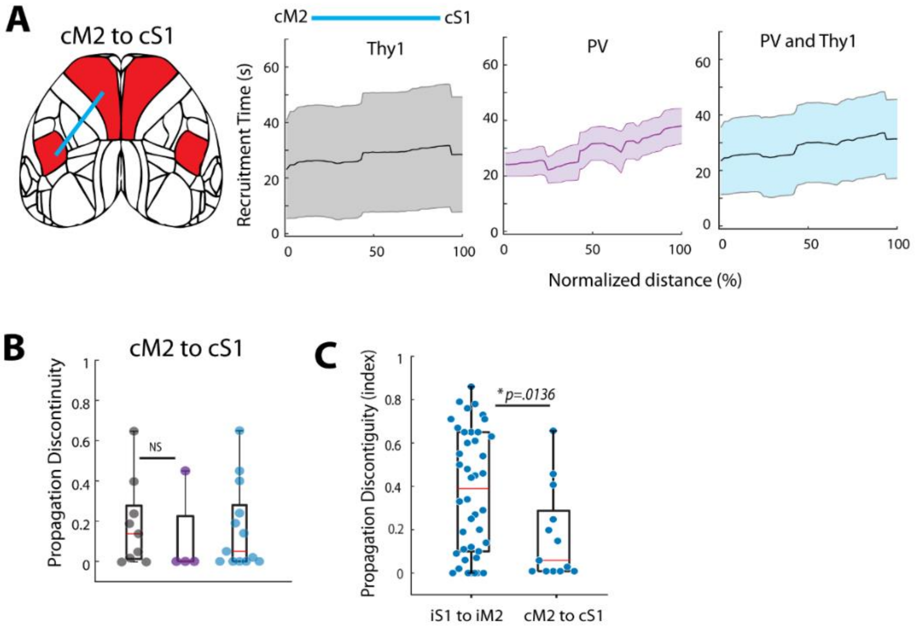


**Supplemental Figure 2.** Propagation discontiguity index calculation in contralateral cortex. A) A direct line of pixels was extracted from propagation maps between iS1 and iM2 using the Bresenham's line algorithm (main Figure 3), here shown for cM2 and cS1. B) Discontiguity calculated between Thy1 and PV animals in seizures that recruited both cM2 and cS1. C) The propagation discontiguity index was significantly higher for iS1-iM2 than for cM2 to cS1 (p=0.013, Wilcoxon rank sum, n=42 iS1-iM2, n=13 cM2-cS1), suggesting that these 4-AP seizures spread through anatomical connections between iS1 and iM2, but then spread more contiguously after crossing the corpus callosum.

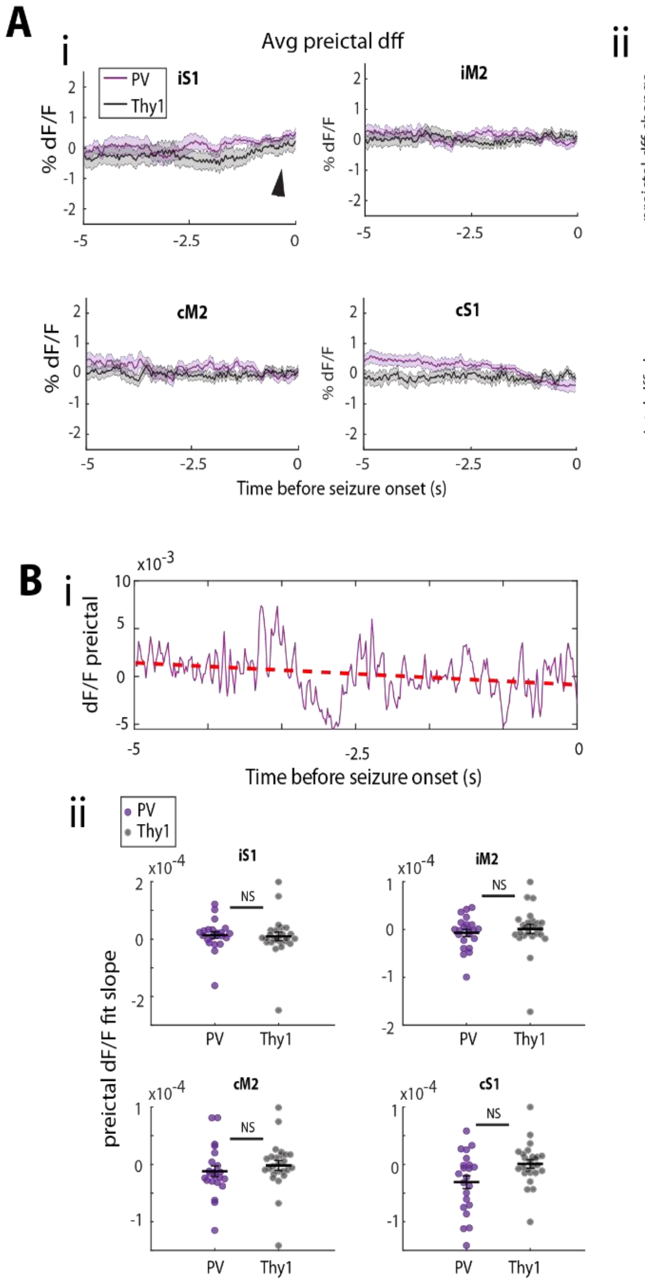


**Supplemental Figure 3.** A) Average preictal dF/F (i) was not different between Thy1 and PV cell types at any S1-M2 site (ii), though a slight increase can be seen at the seizure focus immediately prior to onset (arrow). B) Slope of the GCaMP6f traces (i, sample preictal trace from S1 in a PV-GCaMP6f mouse) was fitted for each preictal period and no difference was seen in Thy1 or PV cell types (ii). Slope of GCaMP traces were calculated in the 5 second preictal period for all seizures. All PV/Thy1 comparisons showed $p>.05$, Wilcoxon rank-sum test n=25 Thy1, n=22 PV seizures.

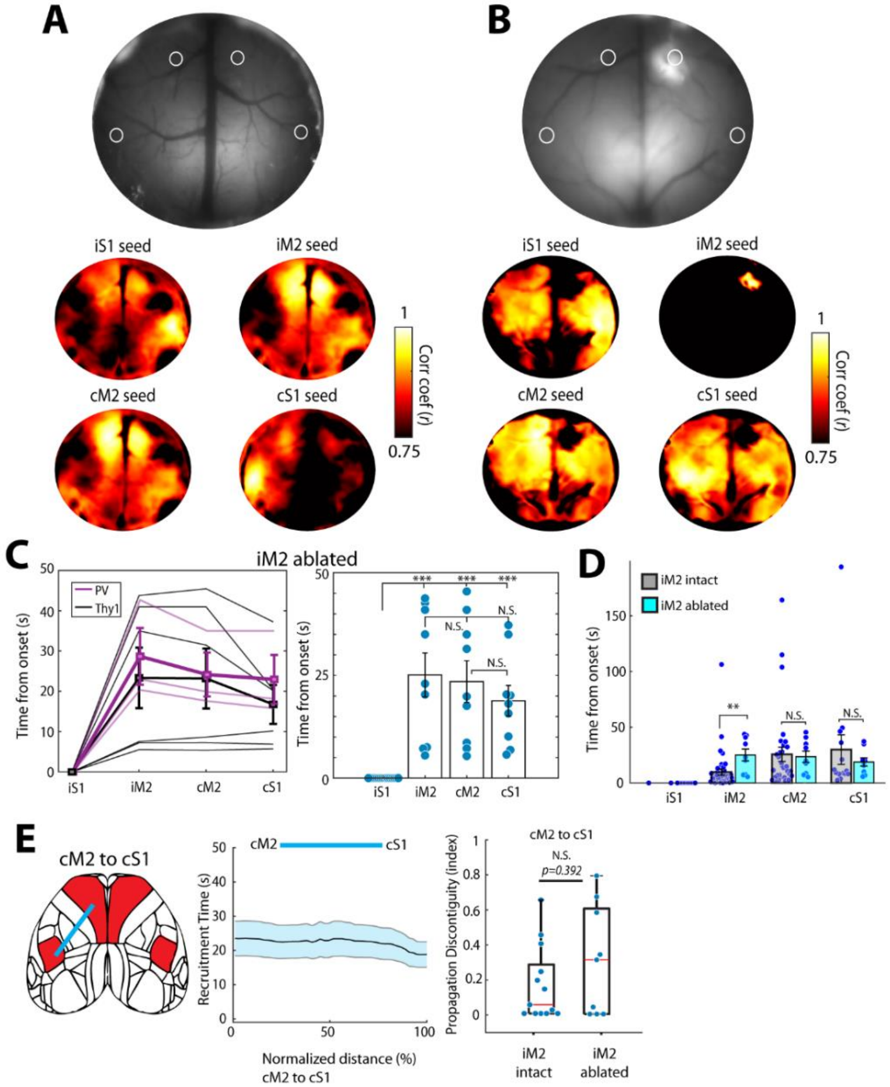


**Supplemental Figure 4.** Ablation of iM2 reduces correlation with the S1-M2 network and affects propagation. A) Cranial window of mouse with iM2 intact, and seed correlation maps showing correlations across the S1-M2 network (bilateral) with seeds placed in each of the nodes. Note the discontinuous connections between iS1, iM2 and cM2. Tthe strongest connections are bidirectionally between iM2 and cM2. B) Same data but for a mouse with right M2 ablated by electrolytic lesioning, showing disruption of the network. Note that a seed placed in iM2 has no correlated pixels and that the absence of iM2 favors contiguous seizure propagation and recruitment in both hemispheres. C) Seizure propagation times between the network nodes following ablation (4 mice, n=16 seizures; *** p<.001, Wilcoxon rank-sum test). D) Seizure onset times compared between iM2 intact and ablated mice. All comparisons were not significant except for iM2, which was ablated. (p=0.0029 for iM2; p=0.095 for cM2, p=0.254 for cS1, Wilcoxon rank-sum test, one tail; n=16 iS1 seizures, n-9 seizures in iM2, cM2, cS1.) E) Discontiguity measurement between cM2 and cS1 shows earlier onset, on average, in cS1, though the discontiguity index was not different between ablated and non-ablated mice (p=.0392, Wilcoxon rank-sum test, n=13 iM2 intact seizures, n=9 iM2-ablated seizures)

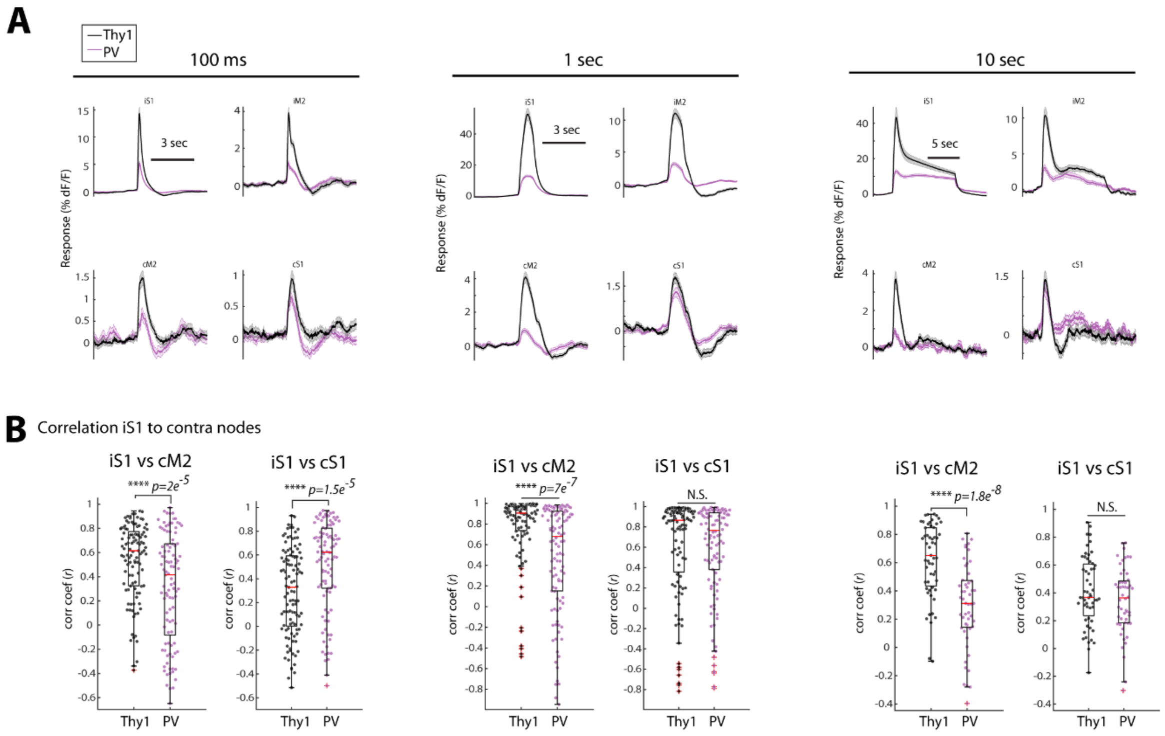


**Supplemental Figure 5.** Changes in fluorescence during electrical stimulation. A) iS1 activity was notably stronger than iM2 across stimulation durations. B) Correlation calculated on the stimulation period between Thy1 and PV. Comparisons are made between Thy1 and PV responses (Wilcoxon rank sum test; 100 ms, 1 sec, and 10 sec trial n=109, 107, 55 Thy1 trials, n=101, 110, 52 PV trials).

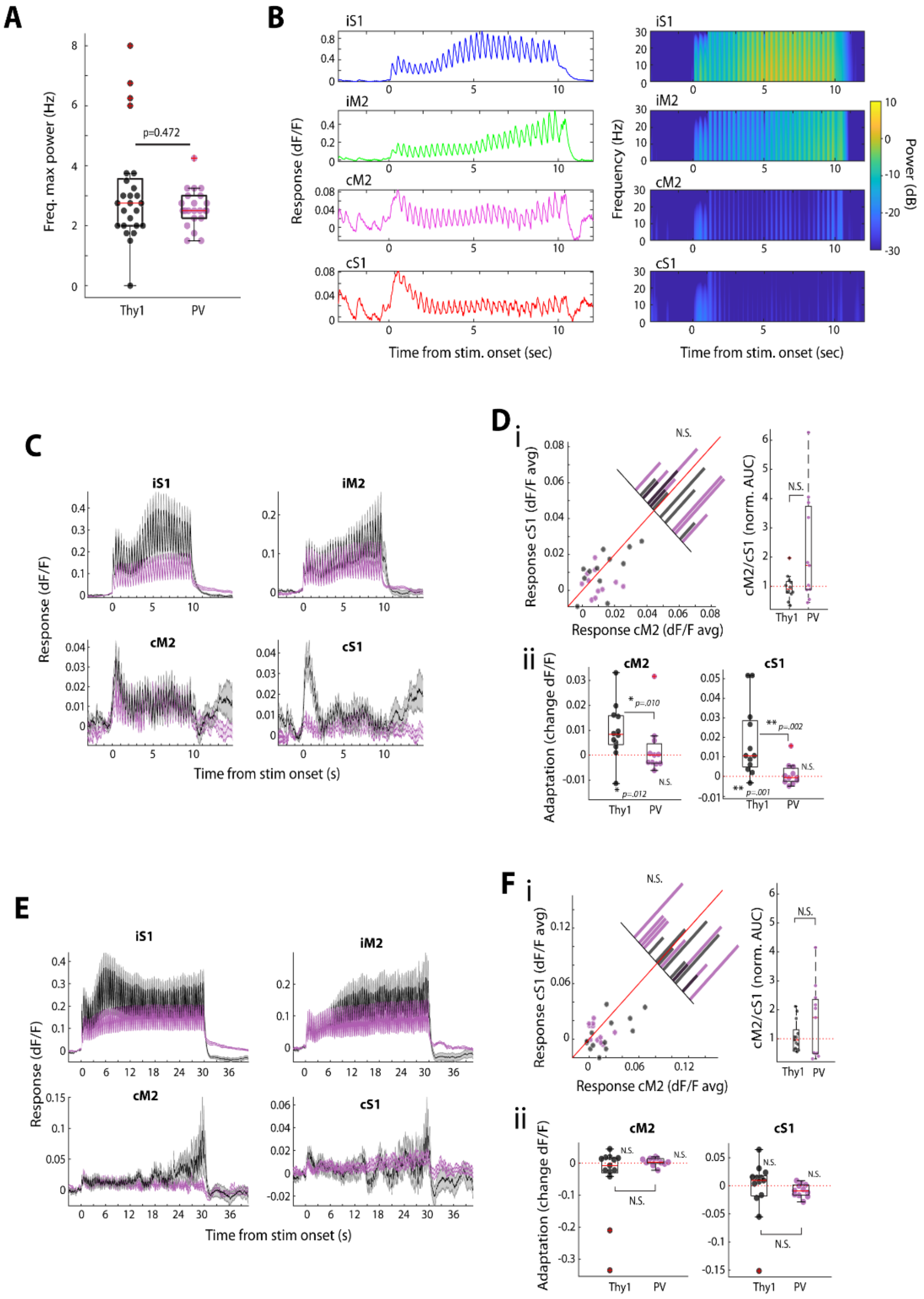

**Supplemental Figure 6.** Ictal-like stimulation. A) The frequency of maximum power was not significantly different between Thy1-GCaMP6f and PV-GCaMP6f animals (p>.05 Wilcoxon rank-sum test) and was ~3 Hz on average. B) Calcium response and power spectra show clear 3 Hz activity recruitment, which facilitates throughout the stimulation period. C) Average responses to seizure-like stimulation in 3 PV, 3 Thy mice (n=12 Thy1 trials, n=11 PV trials). D) i) Activity (dF/F) during 3 Hz stimulation at cS1 and cM2; ii, Adaptation was significant in Thy1 cells at cM2 and cS1, though this is due to an initial response followed by weak activity (see traces in C) throughout the stimulation, which was weaker in PV recordings. E) Same as C for 30 second stimulation trials (n=13 Thy1, n=11 PV). F) Same as D for 30 second stimulation trials. No significant differences were observed between Thy1 and PV in these longer trials.
